# Scaling the Functional Nanopore (FuN) Screen: Systematic Evaluation of Self-Assembling Membrane Peptides and Extension with a K^+^-Responsive Fluorescent Protein Sensor

**DOI:** 10.1101/2022.12.16.520551

**Authors:** Klara Eisenhauer, Wadim Weber, Philipp Kemp, Carolin Gebhardt, Marwan Kaufmann, Noel Tewes, Hanna Zhadanowa, Alesia Tietze, Oliver Rauh, Viktor Stein

## Abstract

The functional analysis of protein nanopores is typically conducted in planar lipid bilayers or liposomes exploiting high-resolution but low-throughput electrical and optical read-outs. Yet, the reconstitution of protein nanopores *in vitro* still constitutes an empiric and low-throughput process. Addressing these limitations, nanopores can now be analyzed using the functional nanopore (FuN) screen exploiting genetically-encoded fluorescent protein sensors that resolve distinct nanopore-dependent Ca^2+^ in- and efflux patterns across the inner membrane of *Escherichia coli*. With a primary proof-of-concept established for the S^21^68 holin, and thereof based recombinant nanopore assemblies, the question arises to what extent alternative nanopores can be analyzed with the FuN screen and to what extent alternative fluorescent protein sensors can be adapted. Focussing on self-assembling membrane peptides, three sets of 13 different nanopores are assessed for their capacity to form nanopores in the context of the FuN screen. Nanopores tested comprise both natural and computationally-designed nanopores. Further, the FuN screen is extended to K^+^-specific fluorescent protein sensors to provide a complementary read-out to Ca^2+^. Finally, a comparison with high-resolution biophysical and electrophysiological studies in planar lipid bilayers provides an experimental benchmark for future studies.

## Introduction

Protein nanopores comprise a versatile class of membrane proteins that are increasingly exploited across a range of biotechnological applications^1^, most prominently DNA sequencing^2^ and more recently proteins^3,4^. Further, nanopores can be modified with receptors turning them into exquisitely sensitive and specific molecular sensors^1^. A key bottleneck concerns our ability to study and engineer nanopores by experimental means which frequently relies on tailored expression and reconstitution protocols in lipid bilayers *in vitro*^5–7^. Addressing these limitations, the functional nanopore (FuN) screen exploits Ca^2+^-responsive fluorescent protein (FP) sensors to resolve the functional properties of a nanopore according to distinct Ca^2+^ in- and efflux patterns across the *E. coli* inner membrane^8^ (**Fig. 1A**).

**Figure 1:**
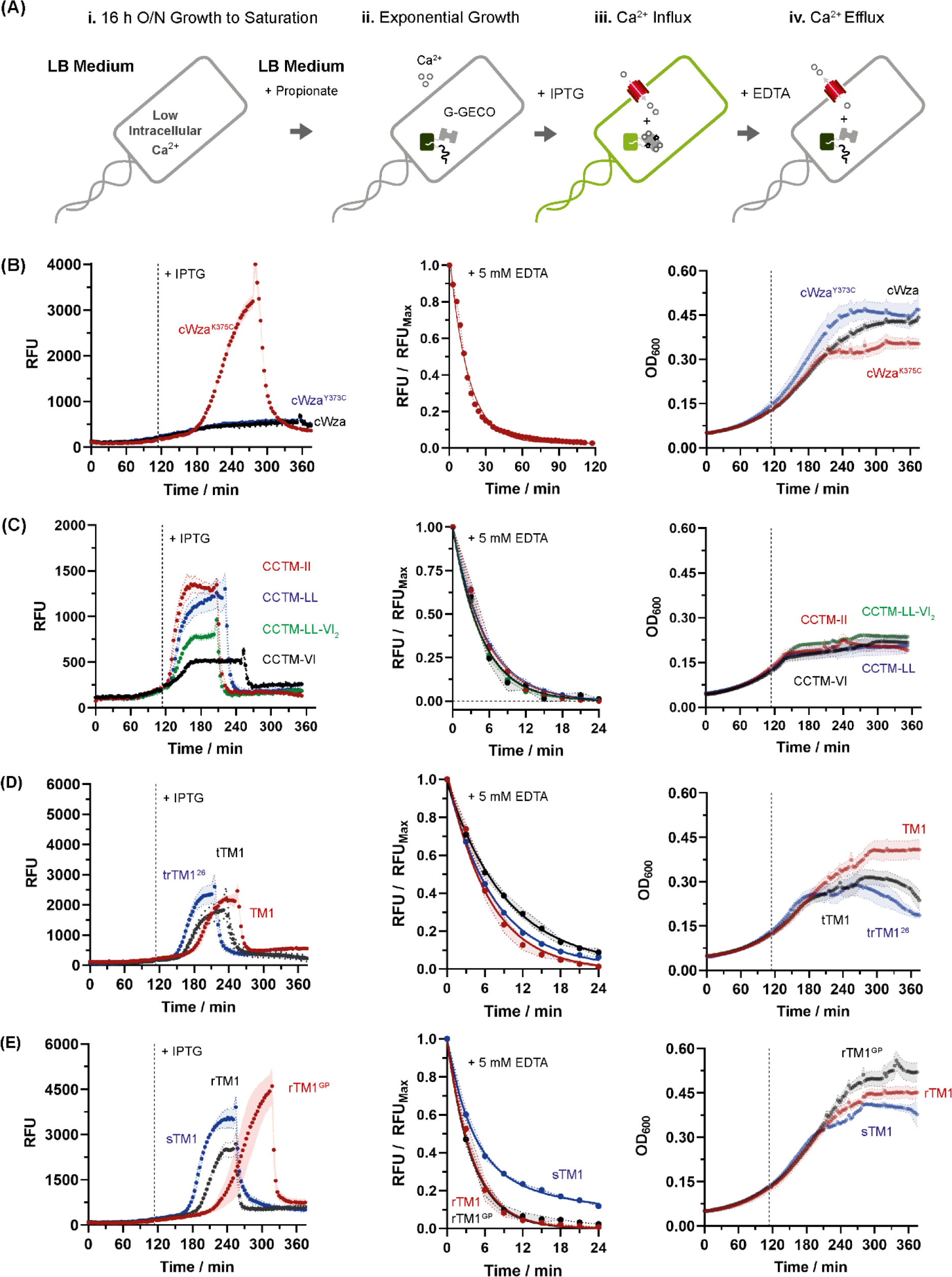
Schematic overview of the FuN screen: **(A)** Ca^2+^-specific FP sensors resolve the functional properties of protein nanopores according to distinct Ca^2+^ in- and effluxes patterns across the inner membrane of *E. coli*; **(B-E)** Summary of kinetic traces used to characterize the functional properties of 13 different nanopores based on the T_½_^IPTG^ and T_½_^EDTA^. The EDTA-dependent decay is shown in a normalized fashion to facilitate comparisons. Nanopores tested include **(B)** a consensus design stabilized transmembrane module of cWza, **(C)** a set of computationally-designed nanopores based on α-helical coiled-coil transmembrane (CCTM) motifs, and **(D, E)** a set of nanopores that have been derived from the first transmembrane (TM1) domain of the Mϕ D29 holin. Nanopores were expressed in the context of the pCtrl2 from the stronger promoter (T7.100.WT) using 0.5 mM IPTG before fluorescence was quenched with 5 mM EDTA. An averaged, time-dependent signal based on three independently picked colonies is shown.

Given its recent implementation and proof-of-concept with the S^21^68 holin and thereof based recombinant assemblies, a question concerns to what extent alternative nanopores can be resolved with the FuN screen. Focussing on self-assembling membrane peptides, a representative set of nanopores was identified from the literature^9–12^ and systematically evaluated using the FuN screen (**Tab. 1, Tab. S1**). Nanopores tested include a consensus design of the pore-forming module of the bacterial transmembrane protein Wza^9^, a set of *de novo* designed nanopores featuring α-helical coiled-coil transmembrane (CCTM) structures^10^, and the first transmembrane (TM1) domain of the Mϕ D29 holin^11,12^. Further, the scalability of the FuN screen principle is demonstrated with a K^+^-responsive FP sensor which now provides a complementary read-out to Ca^2+^ to assess the specificity of nanopores and ion channels reconstituted in the inner membrane of *E. coli*.

**Table 1:** Systematic evaluation of 13 self-assembling membrane peptides in the context of the FuN screen at two different promoter strengths using Ca^2+^-responsive FP sensors.

| Nanopore | T7.100.sRBS |  | T7.100.wt.sRBS |  | Key Features According to Original Publication |
| --- | --- | --- | --- | --- | --- |
|  | T<sub>1/2</sub><sup>IPTG</sup> | T<sub>1/2</sub><sup>EDTA</sup> | T<sub>1/2</sub><sup>IPTG</sup> | T<sub>1/2</sub><sup>EDTA</sup> |  |
| cWza | - | - | - | - | Defined P-, L- and H-states<sup>9</sup> |
| cWza<sup>Y373C</sup> | - | - | - | - | Less stabilized H-state<sup>9</sup> |
| cWza<sup>K375C</sup> | - | - | 107.5 ± 0.5 | 11.3 ± 0.1 | More stabilized H-state<sup>9</sup> |
| CCTM-II | 28.2 ± 0.6 | 3.0 ± 0.1 | 19.9 ± 0.9 | 3.8 ± 0.2 | Large Conductivity<sup>10</sup> |
| CCTM-LL | 39.8 ± 0.6 | 3.3 ± 0.1 | 25.0 ± 1.8 | 3.9 ± 0.2 | Gating \| Monodisperse<sup>10</sup> |
| CCTM-VI | 41.9 ± 5.3 | 2.9 ± 0.2 | 33.2 ± 0.6 | 3.2 ± 0.2 | Stable \| Monodisperse<sup>10</sup> |
| CCTM-LL-VI<sub>2</sub> | 37.9 ± 0.7 | 3.5 ± 0.1 | 27.4 ± 0.6 | 3.2 ± 0.1 | New Recombinant |
| D29 trTM<sup>26</sup> | 84.4 ± 0.7 | 5.3 ± 0.1 | 53.2 ± 0.7 | 5.0 ± 0.1 | Membrane Breaker<sup>11</sup> |
| D29 tTM1 | 129.7 ± 1.0 | 5.9 ± 0.1 | 63.5 ± 1.0 | 6.3 ± 0.2 | Membrane Breaker<sup>11</sup> |
| D29 sTM1 | 138.2 ± 0.6 | 6.7 ± 0.2<sup>§</sup> | 79.6 ± 1.9 | 5.0 ± 0.2<sup>§</sup> | Ion Channel<sup>11</sup> |
| D29 TM1 | - | - | 83.6 ± 3.1 | 4.7 ± 0.3 | Membrane Breaker<sup>11</sup> |
| D29 rTM1 | - | - | 93.4 ± 0.5 | 2.8 ± 0.2 | Ion Channel<sup>11</sup> |
| D29 rTM1<sup>GP</sup> | - | - | 147.4 ± 2.9 | 2.7 ± 0.2 | Ion Channel<sup>11</sup> |

§ Alternatively, a two-phase model yields: T<sub>1/2</sub><sup>EDTA</sup> (Fast): 2.8 ± 0.1; T<sub>1/2</sub><sup>EDTA</sup> (Slow): 20.6 ± 0.9; %-Fast: 0.72 ± 1.1
§ Alternatively, a two-phase model yields: T<sub>1/2</sub><sup>EDTA</sup> (Fast): 3.0 ± 0.2; T<sub>1/2</sub><sup>EDTA</sup> (Slow): 18.3 ± 0.9; %-Fast: 0.59 ± 3.6

## Results

In total, 11 out of 13 nanopores triggered characteristic Ca^2+^-dependent signals upon reconstituting in the *E. coli* inner membrane (**Fig. 1B-E**). In general terms, the fluorescence along with the T_½_^IPTG^ turned out promoter dependent. In fact, for several nanopores – namely, cWza^Y373C^, D29 rTM1^GP^, D29 rTM1 and to some extent D29 TM1 – expression from the stronger promoter turned out critical in generating a sigmoidally-shaped nanopore-dependent curve in the first place (**Fig. S1, Fig. S3**). Conversely, promoter strength did not impact the EDTA-dependent efflux rate, T_½_^EDTA^, as it provides a measure for the functional integrity of a nanopore independent of the underlying transcription, translation and assembly kinetics (**Tab. 1**). Further, a limited number of nanopores (e.g. cWza and cWza^Y373C^) turned out toxic according to plateauing OD_600_s but did not trigger a Ca^2+^-dependent signal in the FuN screen (**Fig. 1B, Fig. S1**).

Further, a comparison with high-resolution biophysical studies recapitulates the formation of 11 out of 13 nanopores tested. In case of a minimal Wza-derived transmembrane module^9^ only a disulphide-stabilized mutant, cWza^K375C^, but neither the primary consensus design, cWza, nor an alternative disulphide stabilized variant, cWza^Y373C^, triggered a fluorescent signal in the FuN screen. While these relative trends potentially reflect the distribution of characteristic closed (C), low (L) and high (H) electrically conductive states (i.e. cWza^K375C^ > cWza > cWza^Y373C^)^9^, it is also conceivable that cWza and cWza^Y373C^ are impermeable to Ca^2+^ or are toxic, but do not effectively assemble in the first place.

Further, a set of *de novo* designed, hexameric α-helical CCTMs^10^ triggered a rapid nanopore-dependent signal with distinct Ca^2+^ in- and efflux patterns (**Fig. 1C, Fig. S2**). Notably, CCTM-II yielded the highest absolute fluorescence along with the fastest T_½_^IPTG^ influx rates correlating with the most conductive states in electrophysiological measurements^10^. In contrast, CCTM-LL and CCTM-VI displayed progressively lower absolute fluorescent signals and proportionately delayed T_½_^IPTG^ influx rates which is conclusive with lesser conductive states observed in electrophysiological measurements^10^. Otherwise, EDTA-dependent efflux rates T_½_^EDTA^ proved highly similar for different CCTM variants suggesting that, at least in the context of the FuN screen, CCTM variants primarily differ in their propensity to assemble into Ca^2+^-permeable nanopores. In addition, a hybrid nanopore featuring one LL and two VI motifs at critical α-helix packing sites yielded intermediate Ca^2+^-dependent in- and efflux rates highlighting substantial plasticity in the underlying CCTM scaffold.

Finally, the FuN screen could recapitulate key features underlying the formation of nanopores by the D29 TM1 holin^11,12^. Previous studies showed that D29 TM1 undergoes a structural transition from β-hairpin to α-helix first triggering local membrane depolarization before facilitating further insertion and assembly of D29 TM1 nanopores to result in complete permeabilization^11,12^. Strikingly, sequence-permutated variants are limited to α-helical conformations and form defined ion channels^11,12^. Notably, the FuN screen recapitulates many of these features as variants with a strong β-hairpin propensity – namely, TM1, tTM1 and trTM1^26^ – triggered an early increase in the Ca^2+^-dependent signal as judged by a T_½_^IPTG^ (**Fig. 1D, Fig. S3**). In contrast, judging by pro-longed T_½_^IPTG^, D29 TM1 variants that are restricted to a α-helical conformation – namely, rTM1 and rTM1^GP^ – need to accumulate to a higher concentration – before triggering a late, but strong increase in fluorescence (**Fig. 1E, Fig. S3**). Conversely, EDTA-dependent T_½_^EDTA^ efflux rates were reduced approximately 2-fold for TM1, tTM1 and trTM1^26^ relative to rTM1 and rTM1^GP^ (**Tab. 1**). This could for once be due to fewer nanopores being synthesized before the inner membrane is permeabilized and adversely impacts their expression. Alternatively, this may arise from the formation of more transient and thus less Ca^2+^ permeable nanopores as has been implicated in prior electrophysiological measurements^12^. Only sTM1 proves an exception as it forms nanopores early while displaying slower EDTA-dependent efflux rates despite a strong α-helical propensity in lipid micelles and ion channel-like properties in electrophysiological measurements^12^. In this regard, one should note that a two-phase exponential decay provides a better fit for EDTA-dependent efflux rates compared to a one-phase model (**Fig. S4**). In fact, in this case, the faster EDTA-dependent efflux rate approximates the efflux rate observed for rTM1 and rTM1^GP^ which potentially implicates two distinct populations of high and low conductive sTM1.

As the FuN screen recapitulates the functional reconstitution of 11 out of 13 self-assembling nanopores, several questions arise. In particular, cWza and cWza^Y373C^ did not yield a nanopore-dependent change in the fluorescent signal (**Fig. 1B**) which could either be due to nanopores not forming in the inner membrane in the first place, or nanopores being impermeable to Ca^2+^. In this context, it must be considered that nanopores were previously characterized with K^+^ *in vitro*^9–12^ while the FuN screen exploits Ca^2+^ as a permeant^8^. To address this discrepancy, the FuN screen was extended with genetic indicators for K^+^ optical imaging (GINKOs). These refer to a new class of intensiometric FP sensors that can detect K^+^ in live cells^13^. Given an apparent K_D_ in the mM range, GINKOs generally saturate at intracellular K^+^ of 200-300 mM. An experimental protocol was thus devised where nanopore reconstitution triggers a K^+^ efflux and thus decrease in fluorescence before rescuing it with 200 mM KCl (**Fig. 2A**).

**Figure 2:**
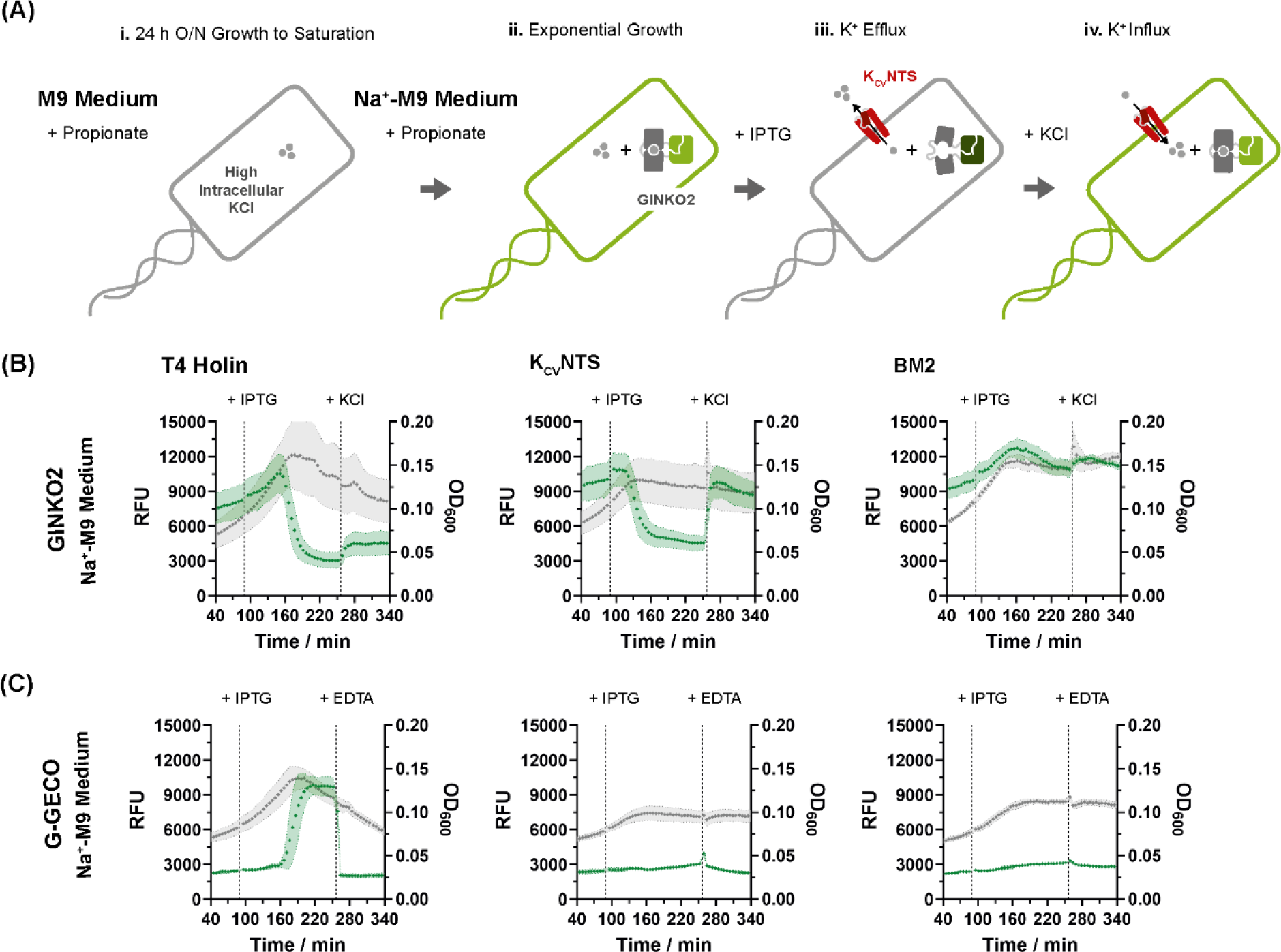
Extending the FuN screen principle to resolve the permeation of K^+^ ions. **(A)** A K^+^-responsive FP sensor, termed GINKO2, resolves distinct nanopore dependent fluxes of K^+^ across the inner membrane of *E. coli*. Given high intracellular K^+^ concentrations, fluorescence first decreases as K^+^ diffuses out of the cell upon formation of a nanopore in the inner membrane but is subsequently rescued with 200 mM KCl; **(B)** Summary of K^+^-dependent fluorescent signals for different nanopores and ion channels expressed in *E. coli* grown in K^+^-deficient Na^+^-M9 medium. The large, lytic T4 holin and the defined K^+^-specific ion channel K_CV_NTS trigger a decay in fluorescence but only K_CV_NTS can be quantitatively rescued with 200 mM KCl. Conversely, the H^+^-specific BM2 ion channel remains impermeable to K^+^; **(C)** Summary of Ca^2+^-dependent fluorescence in Na^+^-M9 medium. The large, lytic,T4 holin triggers an increase in fluorescence which is subsequently quenched with 5 mM EDTA. Conversely, the K^+^-permeable K_CV_NTS and the H^+^-permeable BM2 channel remain impermeable to Ca^2+^.

To this end, *E. coli* were grown in modified K^+^-deficient Na^+^-M9 medium (**Tab. S2**) to generate a K^+^ gradient across the inner membrane that can be effectively resolved across the dynamic range of GINKO2. To this end, GINKO2 had to be pre-expressed for 24 h as its comparatively slow maturation kinetics would otherwise obscure the K^+^-dependent decrease in fluorescence upon formation of a nanopore in the inner membrane. Following protocol optimization, the lytic T4 holin^14^ and K^+^-specific K_CV_NTS^15^ but not the H^+^-specific BM2^16^ triggered a nanopore-dependent decrease in fluorescence (**Fig. 2B**). Crucially, fluorescence could be rescued with 200 mM KCl for K_CV_NTS, but not for the lytic T4 holin. The latter could be attributed to cell lysis triggering an exchange with Na^+^-M9 medium which reduced the K^+^-dependent response approximately 6-fold (**Fig. S5**). Otherwise, the relative nanopore-dependent changes in the Ca^2+^ fluorescent signal remained unchanged for the T4 holin, K_CV_NTS and BM2 ion channel in Na^+^-M9 medium compared to in LB (**Fig. 2C**).

Next, different cWza, CCTM and D29 TM1 variants were tested for their ability to mediate either an efflux of K^+^ or an influx of Ca^2+^ across the inner membrane of *E. coli* grown in Na^+^-M9 medium (**Fig. 3** and **Fig. 4**). In 10 out of 13 cases, an endpoint was reached as the reconstitution of individual nanopores caused the OD_600_ to plateau and trigger distinct K^+^-and Ca^2+^-dependent changes in fluorescence (**Fig. 3** and **Fig. 4**). Conversely, no endpoint was reached for TM1, cWza and cWza^Y373C^ as cells continued to grow albeit more slowly in Na^+^-M9 medium (**Fig. S6**). Further, judging by the plateauing OD_600_ and quantitative signal rescue with 200 mM KCl, none of the newly tested nanopores turned out lytic despite that CCTM-II, CCTM-LL and a select number of TM1 variants were implicated in the formation of larger oligomeric assemblies.

**Figure 3:**
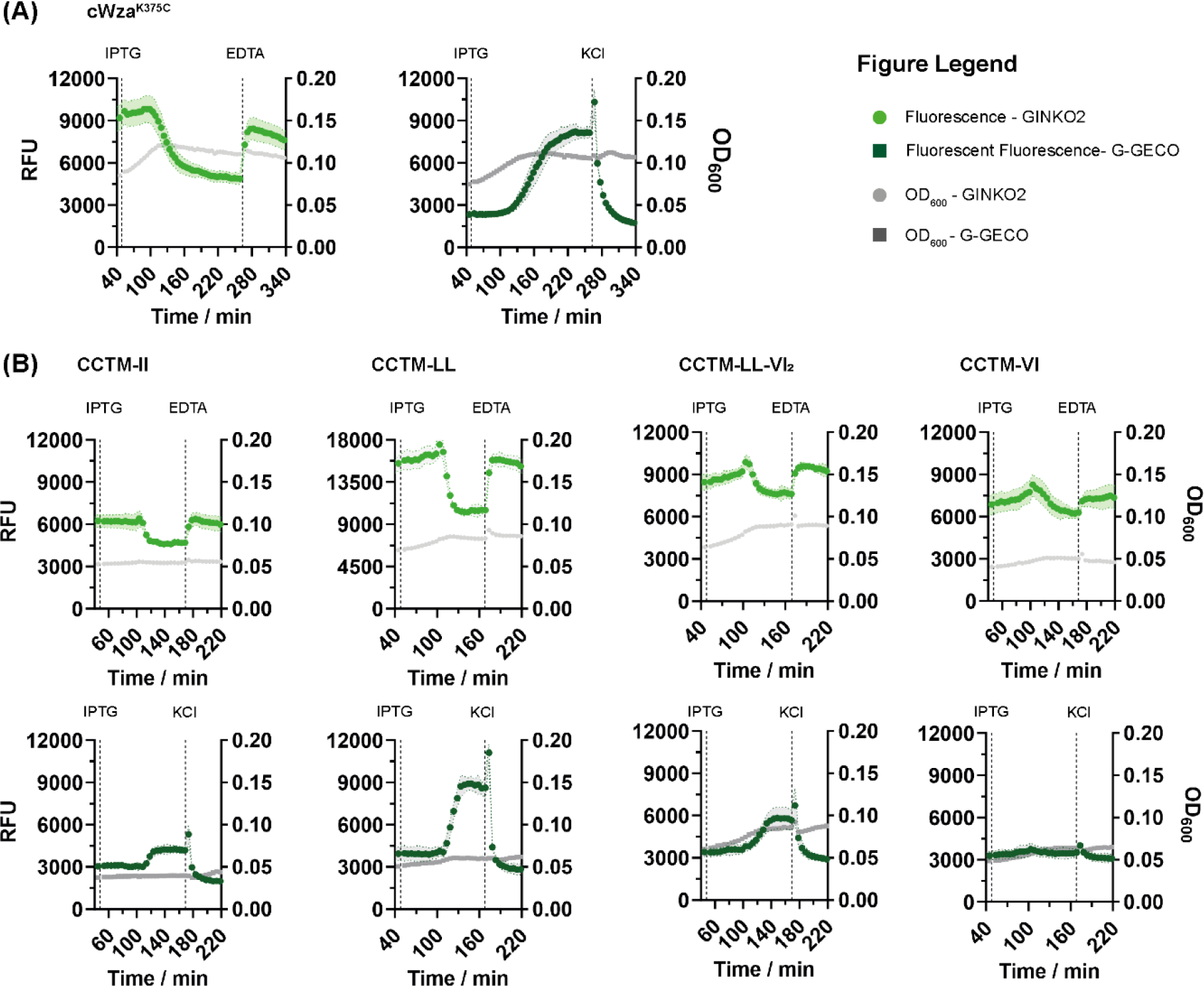
Analysis of cWza and CCTM variants for the permeation of K^+^ and Ca^2+^ following expression in minimal Na^+^-M9 medium. **(A)** cWza^K375C^ is promiscuous for both Ca^2+^ and K^+^ ions; **(B)** CCTM-II, -LL and the new recombinant CCTM-LL-VI_2_ are equally promiscuous for Ca^2+^ and K^+^. Conversely CCTM-VI is permeable to K^+^ but not Ca^2+^. The permeation of K^+^ and Ca^2+^ was resolved either with GINKO2 (light green) or G-GECO (dark green). Nanopores were generally expressed from the strong T7.100.wt.sRBS promoter in *E. coli* grown in Na^+^-M9 medium. The addition of 1 mM IPTG (to induce nanopore expression), and either 5 mM EDTA or 200 mM KCl (to rescue fluorescence) is indicated. Errors are based on three independently picked colonies.

**Figure 4:**
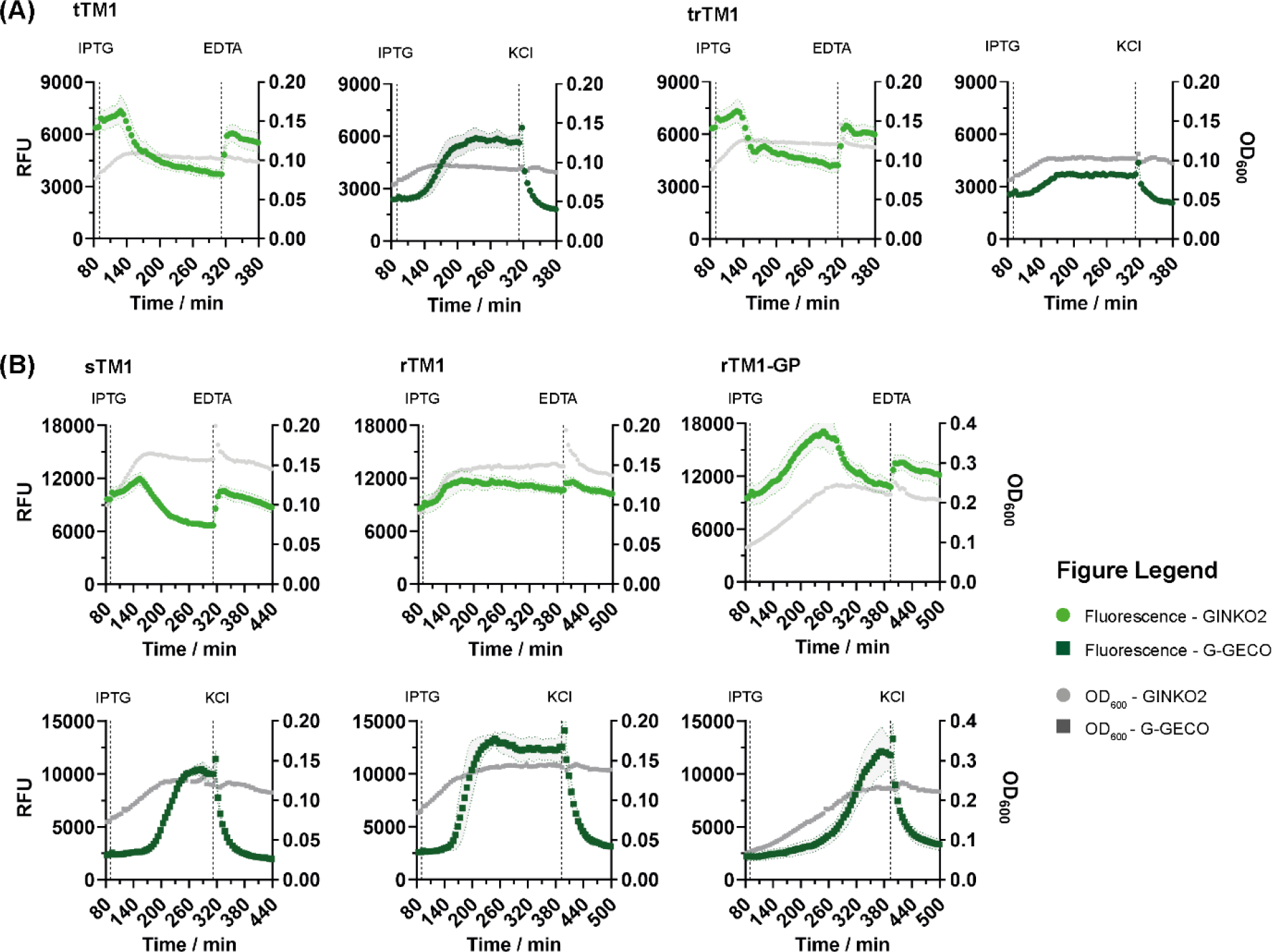
Analysis of D29 TM1 variants for the permeation of K^+^ and Ca^2+^ following expression in minimal Na^+^-M9 medium. **(A)** Membrane breaking β-sheet transitioning tTM1 and trTM1 are promiscuous for both Ca^2+^ and K^+^ ions; **(B)** α-helical, ion channel-like sTM1 and rTM1-GP are equally promiscuous for Ca^2+^ and K^+^. Conversely, rTM1 is permeable to Ca^2+^ but not K^+^ following expression in Na^+^-M9 medium. The permeation of K^+^ and Ca^2+^ was resolved either with GINKO2 (light green) or G-GECO (dark green). Nanopores were expressed from the strong T7.100.wt.sRBS promoter in *E. coli* grown in Na^+^-M9 medium. The addition of 1 mM IPTG (to induce nanopore expression), and either 5 mM EDTA or 200 mM KCl (to rescue fluorescence) is indicated. Errors are based on three independently picked colonies.

Otherwise, the majority of nanopores were permissible to both Ca^2+^ and K^+^. An exception includes CCTM-VI which proved permeable to K^+^ but not Ca^2+^ upon reconstitution in the inner membrane of *E. coli* grown in Na^+^-M9 medium (**Fig. 3B**). The opposite applied to rTM1 which demonstrated a large drop in the Ca^2+^-but not K^+^-dependent fluorescence indicating specificity for Ca^2+^ in the FuN screen (**Fig. 4B**). Otherwise, cWza and cWza^Y373C^ did not reach an endpoint in Na^+^-M9 medium (**Fig. S5**) emulating observations in LB (**Fig. 2B**). Therefore, it remains unclear whether the FuN screen is not sufficiently sensitive to resolve the permeation of Ca^2+^ through either cWza variant, or whether cWza and cWza^Y373C^ do not form nanopores in the first place and their expression – at least in LB medium – is otherwise toxic.

A select number of nanopores were therefore characterized by means of the *in vitro* bilayer technique. In particular, we aimed to discern to what extent different cWza and CCTM variants are permeable to Ca^2+^ especially considering K^+^ was predominantly used as a permeant in previous *in vitro* bilayer measurements^9–12^. To this end, nanopores were chemically synthesized (**Tab. S3** and **S4**) before being applied to vertical DPhPC bilayers and their electrical conductance analyzed in 500 mM CaCl_2_ (which equates to 1 M KCl in previous *in vitro* bilayer measurements^9,10^). Crucially, both cWza^Y373C^ and cWza^K375C^ proved conductive to Ca^2+^ with two populations of conductive states of 0.054 nS and 0.22 nS for cWza^K375C^ (**Fig. 5A**), and 0.09 nS and 0.19 nS for cWza^Y373C^ in 500 mM CaCl_2_ (**Fig. 5B**). This proves both nanopores are generally permeable to Ca^2+^ even though the conductance is decreased approximately 5-fold compared to in 1 M KCl^9^. Further, this suggests discrepancies observed in the FuN screen relate to the ability of individual cWza variants to form nanopores in the *E. coli* inner membrane in the first place.

**Figure 5:**
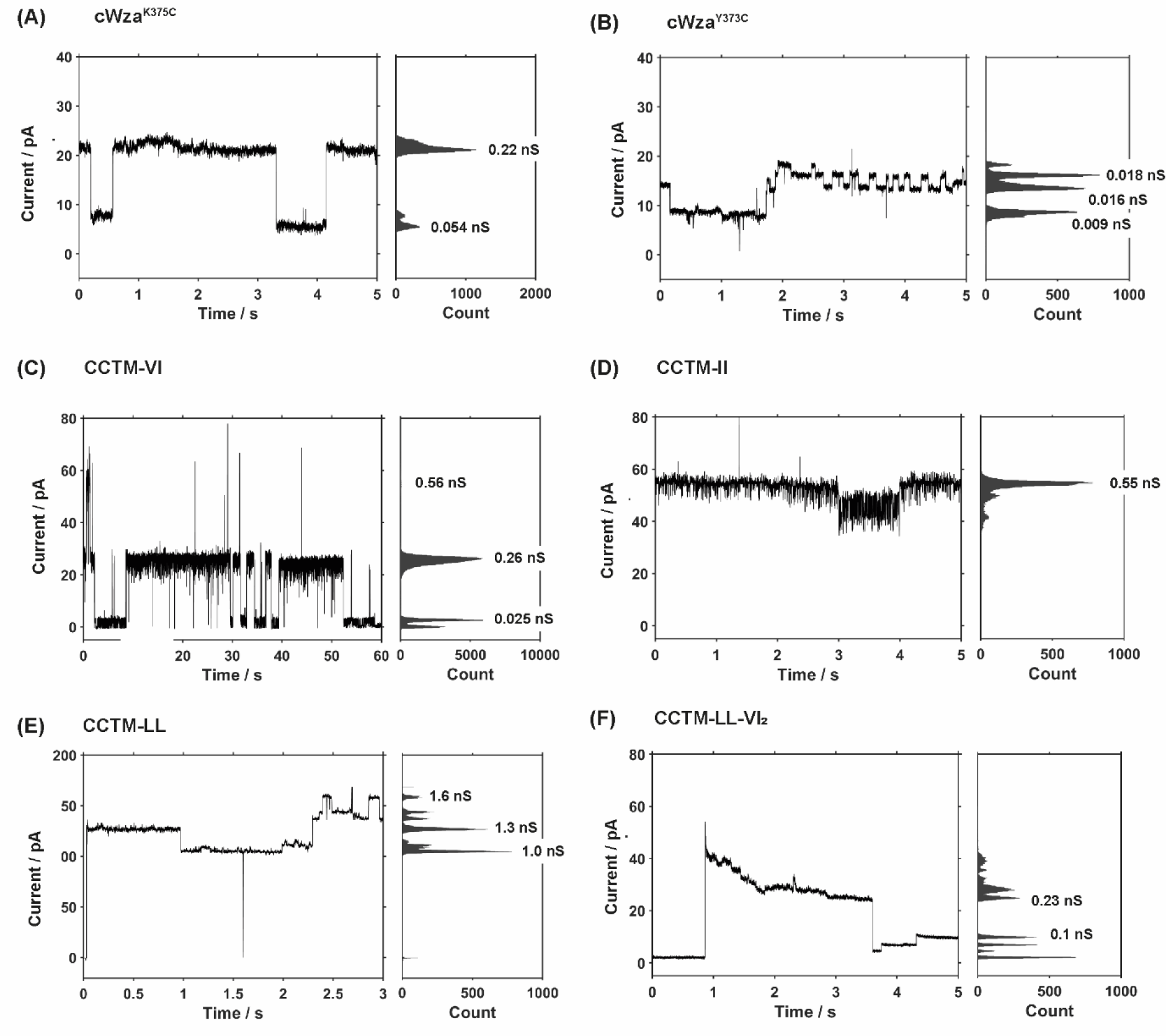
Electrical conductance of a select number of cWza and CCTM variants in DPhPC bilayers. **(A-B)** cWza^K375C^ and cWza^Y373C^ were analyzed in 10 mM HEPES-Ca(OH)_2_, 500 mM CaCl_2_ pH 7.4 at +100 mV, and both conduct Ca^2+^. **(A)** Two distinct states are observed for cWza^K375C^ with a conductance of 0.22 nS and 0.054 nS; **(B)** Similarly, two distinct albeit more heterogenous states are observed for cWza^Y373C^ with a conductance in the range of 0.09 nS and 0.18 nS; **(C-F)** CCTM-VI, CCTM-II, CCTM-LL and CCTM-LL-VI_2_ were analyzed in 10 mM Tris-HCl, 500 mM CaCl_2_ pH 8.0 at +100 mV, and all conduct Ca^2+^. **(C)** In line with previous *in vitro* bilayer measurements, CCTM-VI displays stable ion channel-like gating with a conductance of 0.26 nS; **(D)** CCTM-II displays a conductance of 0.55 nS which either equates to two or one larger nanopore assembly; **(E)** CCTM-LL display multiple heterogenous states with a conductance >1 nS; **(F)** The new recombinant CCTM-LL-VI_2_ variants displays multiple heterogenous states with a steep initial increase which then levels between 0.23 – 0.25 nS and subsequently transitions to a conductance of 0.1 nS.

Similarly, CCTM variants generally proved conductive to Ca^2+^ (**Fig. 5C-F**) underpinning their ability to permeate Ca^2+^ in the FuN screen (**Fig. 1B** and **3B**). In line with previous *in vitro* bilayer measurements^10^, CCTM-VI displayed defined gating and the most defined conductance of 0.26 nS in 500 mM CaCl_2_ (**Fig. 5C**). Please note, this is slightly larger, but still comparable in magnitude relative to previous single channel conductivities of 0.15 nS measured in 1 M KCl^10^. This was followed by CCTM-II with a 2-fold larger conductivity of 0.55 nS in 500 mM CaCl_2_ (**Fig. 5D)** which either equates to two nanopores or one larger nanopore assembly with more than six subunits as has previously been implicated^10^. Further, CCTM-LL displayed large heterogenous but defined conductivities >1 nS (**Fig. 5E**). Finally, the new recombinant CCTM-LL-VI_2_ variant, which combines one heterogenous ‘LL’ and two defined ‘VI’ feature motifs, initially displayed a short-lived peak before adopting more defined conductive states of 0.23 nS – 0.27 nS, and eventually transitioning to a conductance of 0.1 nS −0.14 nS in 500 mM CaCl_2_ (**Fig. 5F** and **Fig. S7**).

## Discussion

To conclude, systematically evaluating the pore-forming propensity of 13 different self-assembling membrane peptides underpins the FuN screen as a complementary approach to high-resolution biophysical studies in reconstituted lipid bilayers. While it lacks the resolution, sensitivity and specificity of electrophysiological measurements^5^, it combines the ease of cellular assays with throughput and kinetic resolution. Further, the scalability of the FuN screen is demonstrated by adapting a K^+^-specific FP sensor. Crucially, the FuN-K^+^ screen implementation can now resolve the permeation of K^+^ ions across nanopores and ion channels paving the way for both foundational studies and more complex engineering efforts such as changing the permeation specificity. Yet, in a limited number of instances, our benchmarking study also highlights both qualitative and quantitative discrepancies compared with previous biophysical and electrophysiological studies. For instance, in case of cWza and cWza^Y373C^, no functional read-out is generated for either Ca^2+^ or K^+^ across any experimental conditions. Further, rTM1 displays no permeability to K^+^ in the FuN screen but defined conductive states in electrophysiological measurements *in vitro*^12^. Finally, CCTM-VI displays the most defined conductive states *in vitro* but the lowest permeability to either Ca^2+^ or K^+^ relative to other CCTM variants in the FuN screen.

To account for these discrepancies, several factors require attention. Firstly, in terms of experimental resolution, the *in vitro* bilayer technique comprises a single molecule technique which can resolve distinct assembly and conformational states on a time scale of ms. In contrast, the FuN screen measures the population averaged permeation properties of nanopores and ion channels inserted in the inner membrane of *E. coli*. The fluorescent signal will therefore be affected by the number of nanopores and its individual permeation properties. Secondly, in terms of delivery, nanopores are applied to a lipid bilayer in purified form and frequently necessitate the careful addition of detergent such as DDM. In contrast, the insertion of nanopores in the inner membrane of *E. coli* occurs autonomously upon recombinant expression which must also mitigate toxicity and functional reconstitution in the uninduced and induced states. Further, the efficiency of insertion along with distinct assembly and potentially conformational states will be influenced by membrane composition. *In vitro*, membranes are highly defined, yet may also feature non-natural lipids such as DPhPC and potentially contain residual detergent and organic solvents. In contrast, membrane composition in *E. coli* is heterogenous and may depend on the strain and distinct physiological states – conversely, it is free of organic solvent residues and detergent. Thirdly, *in vitro* bilayer measurements are conducted under controlled experimental conditions and membrane potentials. In contrast, owing to cellular expression, the FuN screen analyzes nanopores and ion channels against a backdrop of cellular processes. This particularly concerns ion homeostasis which actively maintains K^+^ and Ca^2+^ at high and low intracellular concentrations and needs to be overcome by any nanopore or ion channel to trigger a functional read-out in the FuN screen. In addition, cellular co-factors and chaperons may impact the functional reconstitution of nanopores in the FuN screen. For instance, for the bacterial persister HokB, it has previously been shown that its reversible assembly is regulated by the disulphide oxidizing and reducing enzymes DsbA and DsbC in the periplasm^17^. It is conceivable these account for differences in the experimental read-out of different cWza variants. While cWza^K3735C^ and cWza^Y373C^ both form Ca^2+^ permeable nanopores *in vitro*, cWza^Y3733C^ does distinctly not *in vivo* as judged by a complete lack of either Ca^2+^ or K^+^-dependent fluorescent signals in both rich LB and minimal Na^+^-M9 medium. Access of the mutant Cys to disulphide oxidases may for instance differ as K375C is more surface accessible compared to Y373C and thus potentially more amenable to disulphide-dependent dimerization and subsequent nanopore assembly. Yet, further studies are required to better understand how recombinant expression, cellular processes and especially ion homeostasis quantitatively shape key functional properties of different nanopores such as assembly or permeation specificity in the context of the FuN screen.

Our benchmarking study also highlights the scope for further technological improvement and optimization. Firstly, it is critical for any given nanopore or ion channel to reach an endpoint which equates to a plateauing OD_600_ when nanopores and ion channels have effectively formed in the inner membrane and cause cells stop to divide. When no endpoint is reached, higher expression levels can be achieved by optimising the underlying promoter by enhancing expression and promoting assembly into a functional nanopore. In addition, the accuracy of measurements can be improved by normalizing over the amount of a nanopore, ion channel or FP sensor expressed. For instance, at this stage, it remains unclear to what extent a strong nanopore-dependent fluorescent signal is achieved either through a larger number of number or more permeable nanopores. Normalization can be achieved by co-expressing a nanopore with a FP either as a fusion protein or a bicistronic transcriptional unit. The former is more accurate but risks interfering with the functional properties of a nanopore. Conversely, a bicistronic expression construct has a lesser experimental impact on the functional properties of a nanopore, but only normalizes over transcriptional variability.

In concluding considerations, the FuN screen offers a complementary approach to high-resolution electrophysiological measurements and high-throughput (HTP) screens such as liposome display^6^ or hemolytic assays^18^. Key difference across these experimental platforms concern **(i)** experimental resolution, **(ii)** access to the recombinant route along with throughput, **(iii)** ease of reconstituting any given nanopore in a particular membrane, and **(iv)** the permeant to resolve nanopore and ion channel function. Without doubt electrophysiological measurements provide the most defined experimental conditions and highest experimental resolution as it can discern distinct functional properties with single molecule resolution^5–7^. Yet, the recombinant route also proves most challenging as nanopores need to be purified and stabilized in detergent, yet also affords flexibility to introduce non-natural amino acids^19^. Conversely, liposome display affords genetic encoding and very HTP >10^8-9^ but necessitates cell-free expression and the reconstitution of a nanopore in liposomes *in vitro*. Further, a functional read-out depends on the covalent capture of a synthetic fluorophore which limits the approach to the analysis of large nanopores. Hemolytic assays are also amenable to HTP screening in multiwell-plate or colony-on-plate formats. However, any given nanopore needs to be secreted and released in the extracellular medium and be capable of triggering cell lysis for a functional read-out. Finally, the FuN screen exploits genetically-encoded FP sensors to resolve the permeation of distinct ions and small molecules across a nanopore formed either in the inner^8^ or recently in the outer membrane of *E. coli*^20^. Crucially, genetically-encoded FP sensors afford flexibility in experimental formats with variable throughput and kinetic resolution such as microtitre plates, colony-on-plate and potentially FACS. A limitation concerns the fact that a functional analysis necessitates recombinant expression in *E. coli* and occurs against a backdrop of cellular processes and endogenous ion homeostasis. Conversely, the former also offers opportunities, for instance, to take advantage of cellular co-factors and chaperones to assist with folding and assembly. In any case, with continuing progress and increasing significance in nanopore engineering, it will be intriguing to see what kind of alternative nanopores such as the recently developed SV28^21^ can be resolved and engineered using the FuN screen across different applications in basic research and synthetic biology.

## Supporting Information

Supporting Information includes **(i)** a detailed account of the materials and methods and **(ii)** fluorescent recordings of all self-assembling membrane peptides.

## Acknowledgements

The authors acknowledge support by Felix Herrmann assisting with experimental efforts.

