## Supporting Information for "Scaling the Functional Nanopore (FuN) Screen: Systematic Evaluation of Self-Assembling Membrane Peptides and Extension with a K^+^-Responsive Fluorescent Protein Sensor"

**Title**

### Materials and Methods

*General on Methodology*

Protein nanopores were generally expressed in the context of the pCtrl2 which mitigates any basal non-specific expression and therefore toxicity in the uninduced state[1]. To discern concentration dependent effects, nanopores were systematically expressed under the control of two different T7 regulated promoters that provide a medium (T7.100.sRBS) and a strong level (T7.100.WT.sRBS) of nanopore expression. The propensity of individual membrane peptides to assemble into functional nanopores in the inner membrane of *E. coli* was then systematically quantified in terms of the $T_{½}^{IPTG}$, $T_{½}^{EDTA}$ and the $OD_{600}$. Here, $T_{½}^{IPTG}$ refers to the time needed to reach half the maximum fluorescent signal and provides a measure for an influx of $Ca^{2+}$ following expression of a nanopore with 0.5 mM IPTG. Further, the $T_{½}^{EDTA}$ quantifies the decay in the fluorescent signal associated with an efflux of $Ca^{2+}$ following the addition of 5 mM EDTA. Crucially, the $T_{½}^{EDTA}$ provides a measure for the structural integrity of a protein nanopore independent of the underlying transcription, translation and assembly kinetics. Finally, the $OD_{600}$ informs on the toxicity and the lytic capacity of a protein nanopore as plateauing $OD_{600}$s are associated with the formation of toxic nanopores, and decreasing $OD_{600}$s are associated with the formation of lytic nanopores[1].

*Recombinant DNA Work*

The $Ca^{2+}$-responsive FP sensor, G-GECO[2], was generally expressed under a propionate-inducible promoter based on the pPro24[3,4]. A set of self-assembling membrane peptides (**Tab. S1**) was cloned into the pCtrl2[1] *via* Nde1 and Kpn1 sites under the control of a strong and a medium IPTG-inducible promoter which are denoted as T7.100.WT.sRBS and T7.100.sRBS. The coding DNA were generally synthetized through a commercial provider (Sigma Aldrich or Integrated DNA Technologies).

**Table S1**: Amino acid sequences of the 13 self-assembling membrane peptides assessed in the FuN screen

| Nanopore | Sequence | Reference |
|---|---|---|
| cWza | APLVRWNRVISQLVPTITGVHDLTETVRYIKTWPN | 5 |
| cWza$^{Y373C}$ | APLVRWNRVISQLVPTITGVHDLTETVRCIKTWPN | 5 |
| cWza$^{K375C}$ | APLVRWNRVISQLVPTITGVHDLTETVRYICTWPN | 5 |
| CCTM-II | KKKKGSGISAWATIISALATIISALATIISAWATVG | 6 |
| CCTM-LL | KKKKGSGISAWATLLSALATLLSALATLLSAWATVG | 6 |
| CCTM-VI | KKKKGSGISAWATVISALATVISALATVISAWATVG | 6 |
| CCTM-LL-$VI_2$ | KKKKGSGISAWATLLSALATVISALATVISAWATVG | New Recombinant |
| D29 TM1 | KIRETLYYVGTLVPGILGIALIWGGIDA | 7 |
| D29 tTM1 | ADIETLYYVGTLVPGILGIALIWGGRIK | 7 |
| D2 trTM1$^{26}$ | IETLYYVGTLVPGILGIALIWGGRIK | 7 |
| D29 sTM1 | ADIGGWIYVGTLVPGILGIALYLTERIK | 7 |
| D29 rTM1 | ADIGGWILAIGLIPGVLTGVYYLTERIK | 7 |
| D29 rTM1$^{GP}$ | ADIGGWILAIGLIGPVLTGVYYLTERIK | 7 |

*Experimental Characterization of Nanopores by means of the FuN Screen*

The functional nanopore (FuN) screen was conducted according to previously established protocols[1]. Briefly, plasmids coding for the 13 different self-assembling membrane peptides were co-transformed with the $Ca^{2+}$-specific FP sensor G-GECO into BL21(DE3) and plated on LB agar plates supplemented with 100 µg/mL ampicillin (AMP) and 25 µg/mL chloramphenicol (CHL). A total volume of 300 µL lysogeny broth (LB) medium (+100 µg/mL AMP; +25 µg mg/mL CHL) was then inoculated with a single colony and grown overnight in a 96 deep-well microtitre plate at 37 ˚C and 1,050 rpm shaking. The next day, 3.5 µL of the saturated *E. coli* suspension was added to 196.5 µL LB medium (+25 mM sodium propionate pH 8.0; 100 µg/mL AMP; 25 µg/mL CHL) in transparent bottoms 96-well black microtitre plates. Plates were then placed and incubated at 30 ˚C in a microtiter plate reader (TECAN Spark) while the fluorescent signal (Ex. 480 ± 10 nm; Em. 525 ± 10 nm) and the $OD_{600}$ were recorded at 3 min intervals. Note, the $OD_{600}$ indicated in **Fig. 1** and **Fig. S1 - S3** refers to the non-adjusted $OD_{600}$ measured by the microtitre plate reader (TECAN Spark) for a sample volume of 200 µL and needs to be multiplied by a factor of approx. 2 for the $OD_{600}$ per cm. At 114 min, nanopore expression was initiated with 0.5 mM IPTG and the $Ca^{2+}$-dependent fluorescent signal along with the $OD_{600}$ continued to be monitored. To quantify the kinetics of self-assembling membrane peptides, the sigmoidal fluorescent signal curve was fit to **Eq. 1** by non-linear regression and the propensity to form functional nanopores quantified in terms of the time needed to reach half the maximum signal $T_{½}^{ITPG}$.

**Eq. 1**

$$Y_{(x)} = Bottom + \frac{Top}{1 + e^{\frac{-(x - T_{1/2})}{Slope}}}$$

To resolve the functional properties of self-assembling membrane peptides independent of the underlying expression and assembly kinetics, 5 mM EDTA was added following saturation of the fluorescent signal. To provide a quantitative measure of EDTA-dependent quench rates, the decaying fluorescent signal was fit to **Eq. 2** by non-linear regression. The half-life associated with the decaying fluorescent signal $T_{½}^{EDTA}$ = ln (2) / k was then used as a measure for the functional properties of a nanopore.

**Eq. 2**

$$Y_{(x)} = (Y_0 - Plateau) \times e^{(-k \times x)} + Plateau$$

Alternatively, the decaying fluorescent signal can be fit to **Eq. 3** by non-linear regression to accommodate for a two-phase exponential decay.

**Eq. 3**

$$Y_{(x)} = f_{fast} \times (Y_0 - Plateau) \times e^{(-k_{fast} \times x)} + f_{slow} \times (Y_0 - Plateau) \times e^{(-k_{slow} \times x)} + Plateau$$

*Experimental Characterization of Nanopores by means of the FuN-$K^+$ Screen*

To resolve the nanopore-dependent permeation of $K^+$ across the inner membrane of *E. coli*, expression constructs coding for different nanopores, ion channels and GINKO2 were transformed into *Escherichia coli* BL21 (DE3) and first plated on LB agar plates supplemented with 25 µg/mL chloramphenicol (CHL) and 50 µg/mL kanamycin (KAN). Then, single colonies were used to inoculate 500 µL M9 medium (+ 25 µg/mL CHL and 50 µg/mL KAN) in 96-deep-well plates. Cultures were then grown for 24 h to saturation at 30 °C under vigorous shaking at 900 rpm. The following day 8 µL cell suspension was transferred into 200 µL minimal $Na^+$-M9 medium (+ 25 µg/mL CHL, 50 µg/mL KAN and 50 mM sodium propionate) in black 96-well plates with transparent bottoms. In $Na^+$-M9 medium, $K^+$ was replaced with $Na^+$ while carry over from the overnight culture resulted in approx. 1.5 mM $K^+$ upon re-inoculation. Cultures were grown in a TECAN microtitre plate reader at 30 °C while the $OD_{600}$ and the developing fluorescent signal (Ex. 480 ± 10 nm; Em. 525 ± 10 nm) was monitored at 5 min intervals. The expression of individual nanopores and ion channels was induced after approximately 2 - 3 h with 1 mM IPTG at the time points indicated when cells reached an $OD_{600}$ of approx. 0.1 – 0.2. Note, the $OD_{600}$ indicated in **Fig. 2 - 4** refers to the non-adjusted $OD_{600}$ measured by the microtitre plate reader (TECAN Spark) for a sample volume of 200 µL and needs to be multiplied by a factor of approx. 2 for the $OD_{600}$ per cm. Once the $OD_{600}$ and the fluorescent signal associated with a nanopore-dependent efflux of $K^+$ stabilized, the fluorescent signal was rescued with 200 mM KCl. The $Ca^{2+}$ permeability in $Na^+$-M9 medium was generally measured in parallel on the same 96-well plate in independently transformed *E. coli* with the G-GECO expression construct. The composition of M9 and $Na^+$-M9 media used in this study is outlined in **Table S2**.

**Table S2.** Summary of composition for adapted M9 and $Na^+$-M9 medium

| Component | Ingredients |
|---|---|
| M9 medium | 1× M9 salts, 0.2% arabinose, 0.05% glucose, 0.1% casein hydrolysate, 1× trace elements, 1 mM $MgSO_4$, 100 µM $CaCl_2$, 0.1% casamino acids (w/v) and 1 µM $FeSO_4$ |
| 10× M9 salts | 420 mM $Na_2HPO_4$, 220 mM $KH_2PO_4$; 90 mM NaCl and 50 mM $NH_4Cl$ |
| 1000× trace elements | 3 mM $(NH_4)_6Mo_7O_{24}$, 400 mM $H_3BO_3$, 30 mM $CoCl_2$, 10 mM $CuSO_4$, 80 mM $MnCl_2$ and 10 mM $ZnSO_4$ |
| $Na^+$-M9 medium | 1× $Na^+$-M9 salts, 0.2% arabinose, 0.05% glucose, 0.1% casein hydrolysate, 1× trace elements, 1 mM MgSO4, 100 µM $CaCl_2$, 0.1% casamino acids (w/v) and 1 µM FeSO4 |
| 10× $Na^+$-M9 salts | 420 mM $Na_2HPO_4$, 220 mM $NaH_2PO_4$, 90 mM NaCl, 50 mM $NH_4Cl$ |

*Chemical Synthesis of cWza and CCTM Membrane Peptides*

The amino acid derivatives used in the study were purchased from Novabiochem, Sigma Aldrich, and Carbolutions and were used without further purification. Intavis® rink amide resin was used as the solid support for the synthesis of the peptides. All solvents and reagents were purchased from commercial sources and used as received.

**Table S3.** List of peptides

| | Name | Sequence | Crude | Purified | Yield |
|---|---|---|---|---|---|
| P1 | cWzaY373C | APLVRWNRVISQLVPTITGVHDLTETVRCIKTWPN | 132.1 mg | 19.1 mg | 15.7% |
| P2 | cWzaK375C | APLVRWNRVISQLVPTITGVHDLTETVRYICTWPN | 118.5 mg | 9.4 mg | 13.4% |
| P3 | CCTM-II | KKKKGSGISAWATIISALATIISALATIISAWATVG | 88.7 mg | 2.7 mg | 4.1% |
| P4 | CCTM-LL | KKKKGSGISAWATLLSALATLLSALATLLSAWATVG | 98.6 mg | 11.3 mg | 3.6% |
| P5 | CCTM-VI | KKKKGSGISAWATVISALATVISALATVISAWATVG | 78.1 mg | 3.46 mg | 3.4% |
| P6 | CCTM-LL-VI2 | KKKKGSGISAWATLLSALATVISALATVISAWATVG | 139.6 mg | 8.3 mg | 5.2% |

Automated Fmoc-SPPS was carried out on an Intavis® MultiPep CF synthesizer. The MultiPep CF one-column module supported by UV monitoring in real-time was used for the synthesis of peptides P1, P2 and P6; peptides P3, P4 and P5 were synthesized using parallel 8-column module (**Table S3**). A standard Fmoc protocol was applied. The synthesis was performed at 100 µmol scale for the peptides P1, P2 and P6, and 50 µmol scale for P3, P4 and P5. Intavis® rink amide resin (0.46 mmol/g) was swelled in 5 ml of dimethylformamide (DMF) for 40 min before the synthesis.

**Synthesis using CF one-column module:** Coupling reactions (double coupling, 40 min) were performed with Fmoc-amino acids (0.5 M, 6 equiv.), activated with 1-[bis(Dimethylamin)methylen]-1H-1,2,3-triazol[4,5-b]pyridinium-3-oxid-hexafluorophosphat (HATU, 0.5 M, 5.93 equiv.) and N-Methylmorpholine (NMM, 0.5 M, 12.00 equiv.) in DMF. For Fmoc deprotection 25% (v/v) piperidine solution in DMF was applied. All deprotection and coupling steps were followed by intensive washing steps with DMF (5 times).

**Synthesis using C8 multi-column module:** Coupling reactions (double coupling, 30 min) were performed with Fmoc-amino acids (0.5 M, 4 equiv.), activated with 2-(1H-Benzotriazole-1-yl)-1,1,3,3-tetramethyluronium hexafluorophosphate (HBTU, 0.5 M, 3.95 equiv.) and N-Methylmorpholine (NMM, 0.5 M, 8.00 equiv.) in DMF. For Fmoc deprotection 20% (v/v) piperidine solution in DMF was applied. All deprotection and coupling steps were followed by intensive washing steps with DMF (5 times).

After the synthesis was completed, all peptides were washed with dichloromethane (DCM) and dried overnight. Peptide cleavage and deprotection were accomplished in a mixture of 95.5% TFA (trifluoroacetic acid), 1% MiliQ $H_2O$, 2.5% 1,2-Ethanedithiol (EDT), and 1% triisopropylsilane (TIPS) for 3 h at room temperature. For 100 mg resin, 3 ml cleavage solution was applied. The crude peptides were precipitated in cold diethyl ether, centrifuged, and washed at least four times with diethyl ether prior to freeze-drying from 60% acetonitrile/water. cWza peptides P1 and P2 were soluble in this mixture; however, CCTM peptides P3, P4, P5 and P6 could not be dissolved in lyo-friendly solvents and were lyophilized as a suspension.

**Solubilization and purification:** Solubilization and purification of P1-P6 are summarized in the **Table S4**. Purification was performed *via* preparative RP-HPLC. Preparative RP-HPLC was performed by Waters$^{TM}$ 600-MS System Controller and Waters$^{TM}$ 2996 Photodiode Array Detector. Flow rate was 8.0 ml/min. The peaks were detected at 220 nm. After purification, collected fractions were combined, freeze-dried and stored at -30°C. The synthesis of all peptides was successful which was confirmed by mass spectroscopy. All peptides were identified by LC-ESI-MS. All products showed the correct molecular weight and isotopic pattern which was in line with the simulated pattern. LC-ESI-MS analysis was performed by using a Waters Q-TOF mass spectrometer. Mass spectra were utilized in positive ion mode over an m/z range of 50–2000.

**Table S4.** Solubilization and purification of peptides

| | Name | Column | Inj. | Gradient and eluents | RT,min |
|---|---|---|---|---|---|
| P1 | cWzaY373C | ISAsphere® 100-10 C4 (PR) (250 x 20 mm) | $H_2O$/AN 1:1 | A($H_2O$):B(ACN) +0.1% TFA, RT 30% to 45% B over 100 min | 46.7 |
| P2 | cWzaK375C | ISAsphere® 100-10 C4 (PR) (250 x 20 mm) | $H_2O$/AN 1:1 | A($H_2O$):B(ACN) +0.1% TFA, RT 38% to 43% B over 100 min | 49.9 |
| P3 | CCTM-II | ISAsphere® 100-5 Si (PR) HILIC (250 x 20 mm) | TFE | A($H_2O$):B(ACN) +0.1% TFA, RT 95% to 70% B over 60 min | 30.0-38.0 |
| P4 | CCTM-LL | ISAsphere® 100-10 C4 (PR) (250 x 20 mm) | HFIP | A($H_2O$):B(iPrOH) +0.1% TFA, 60°C 30% to 100% B over 60 min | 13.0-17.0 |
| P5 | CCTM-VI | ISAsphere® 100-10 C4 (PR) (250 x 20 mm) | HFIP | A($H_2O$):B(iPrOH) +0.1% TFA, 60°C 60% to 100% B over 60 min | 6.0-8.0 |
| P6 | CCTM-LL-VI2 | ISAsphere® 100-5 Si (PR) HILIC (250 x 20 mm) | TFE | A($H_2O$):B(ACN) +0.1% TFA, RT 95% to 70% B over 60 min | 31.0-37.0 |

*Electrophysiological Characterization of cWza and CCTM variants for $Ca^{2+}$ Conductance*

Individual CCTM and cWza variants were characterized at room temperature in a vertical lipid bilayer according to a previously described set up[8]. To this end, chambers were connected to Ag/AgCl electrodes to the head-stage of a patch clamp amplifier L/M-EPC-7 (List-Medical). Membrane potentials were generally referenced to the *cis* compartment. Current traces were filtered at 1 kHz and digitized with a sampling frequency of 5 kHz by an A/D-converter LIH 1600 (HEKA Elektronik). For the analysis of cWza variants, both chambers were filled with 2.5 mL HEPES-based electrolyte (10 mM HEPES-$Ca(OH)_2$, 500 mM $CaCl_2$ pH 7.4). For the analysis of CCTM variants, both chambers were filled with 2.5 mL Tris-based electrolyte (10 mM Tris-HCl, 500 mM $CaCl_2$ pH 8.0). Lipid bilayers were then formed using 1,2-diphytanoyl-sn-glycero-3-phosphocholine (DPhPC) using the air bubble technique[8]. Prior to application to the lipid bilayer, membrane peptides were generally dissolved in DMSO at a concentration of 5-10 mM and then diluted approximately 10-fold in the HEPES-based electrolyte supplemented with 0.05% DDM. Shortly before application to the lipid bilayer, cWza and CCTM peptides were further diluted approximately 50-100-fold in either the HEPES- or Tris-based electrolyte supplemented in 0.005% DDM. Approximately 1-2 µL of the resultant membrane peptide solution was then immediately applied in the vicinity of the lipid bilayer in the *trans* compartment using a Hamilton syringe. The assembly of individual nanopores and their conductance was generally characterized at a constant membrane voltage of +100 mV.

### Supplementary Results

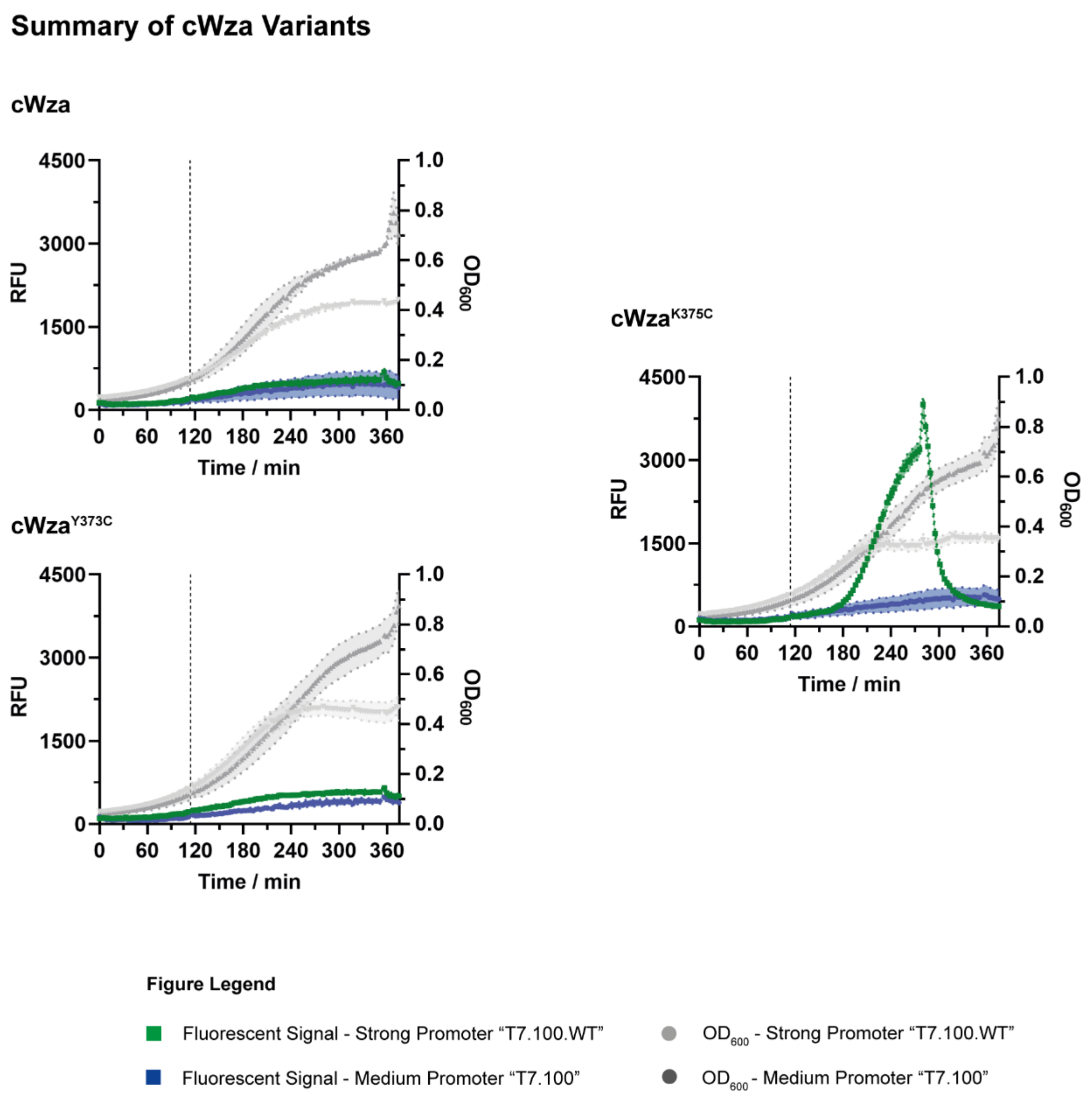

**Figure S1**: FuN screen analysis of cWza variants in LB medium. The expression of individual nanopores was achieved under the control of two different promoters denoted as T7.100.sRBS and T7.100.WT.sRBS that provide medium and strong levels of nanopore expression denoted in green and blue, respectively. Similarly, the course of the $OD_{600}$ is denoted in light and dark grey. Nanopore expression was induced with 0.5 mM IPTG at approximately 114 min and the $Ca^{2+}$-dependent fluorescent signal quenched with 5 mM EDTA.

**Summary of CCTM Variants**

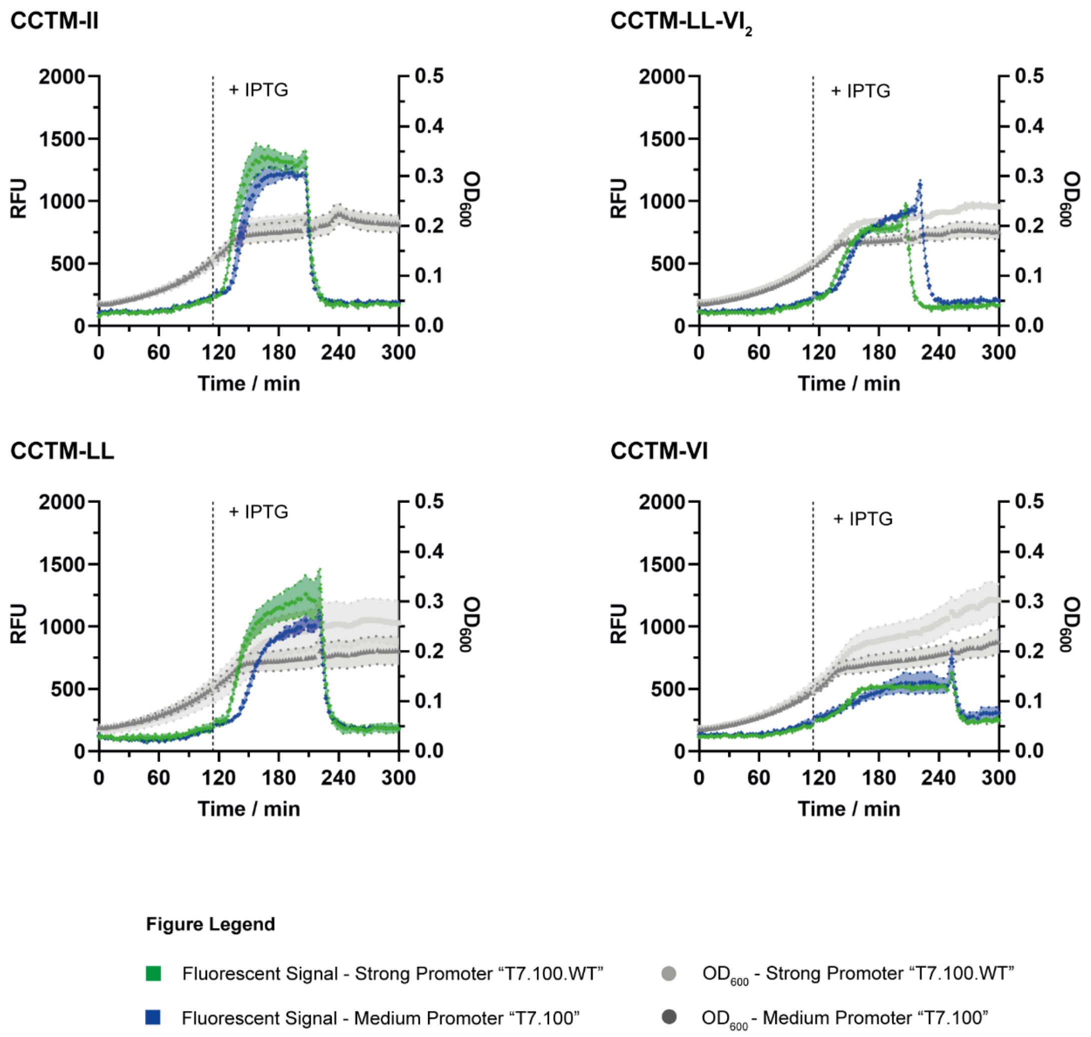

**Figure S2**: FuN screen analysis of CCTM variants in LB medium. The expression of individual nanopores was achieved under the control of two different promoters denoted as T7.100 and T7.100.WT.sRBS that provide medium and strong levels of nanopore expression denoted in green and blue, respectively. Similarly, the course of the $OD_{600}$ is denoted in light and dark grey. Nanopore expression was induced with 0.5 mM IPTG at approximately 114 min and the $Ca^{2+}$-dependent fluorescent signal quenched with 5 mM EDTA.

### Summary of MΦ D29 TM1 Variants

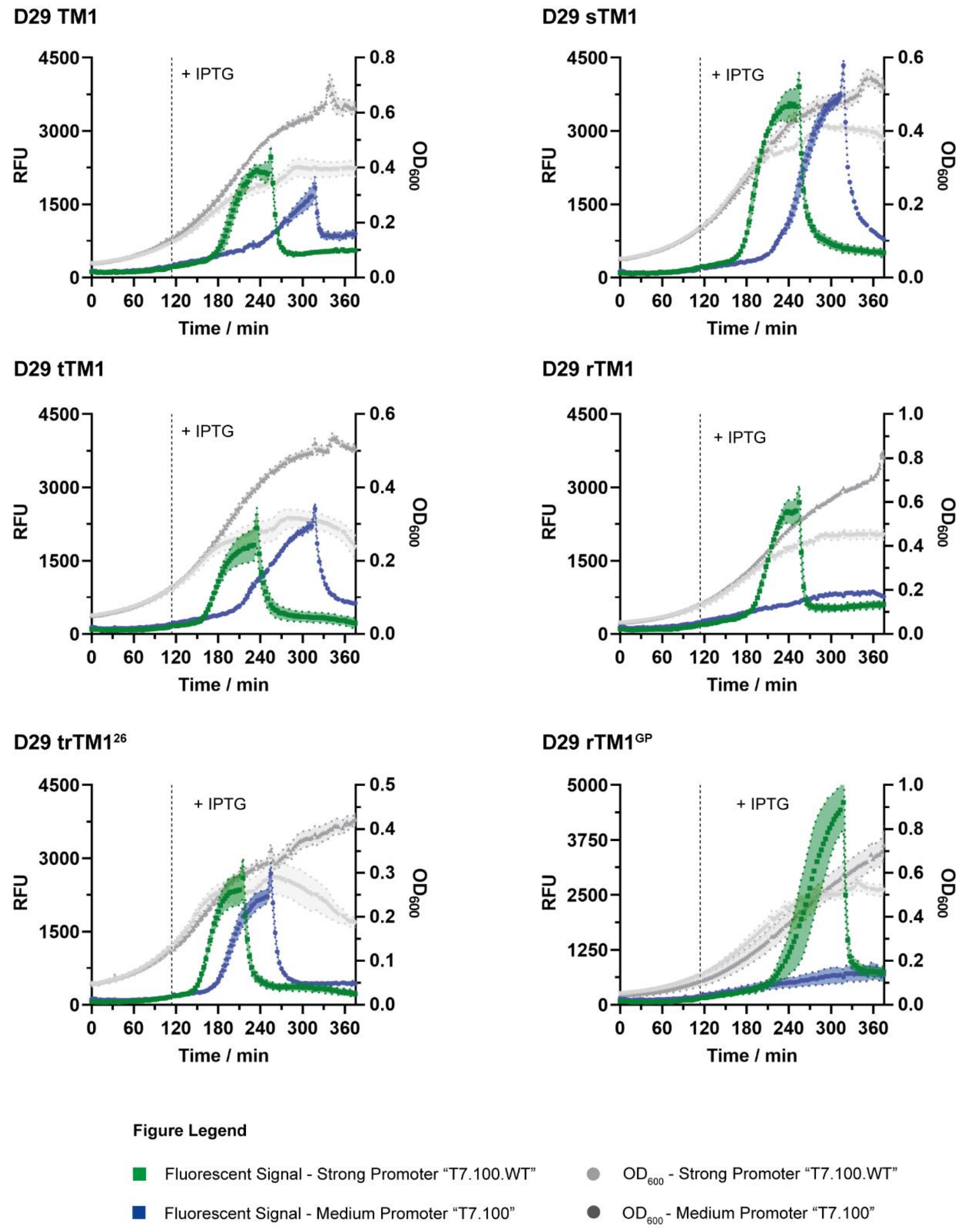

**Figure S3:** FuN screen analysis of D29 TM1 variants in LB medium. The expression of individual nanopores was achieved under the control of two different promoters denoted as T7.100.sRBS and T7.100.WT.sRBS that provide medium and strong levels of nanopore expression denoted in green and blue, respectively. Similarly, the course of the $OD_{600}$ is denoted in light and dark grey. Nanopore expression was induced with 0.5 mM IPTG at approximately 114 min and the $Ca^{2+}$-dependent fluorescent signal quenched with 5 mM EDTA.

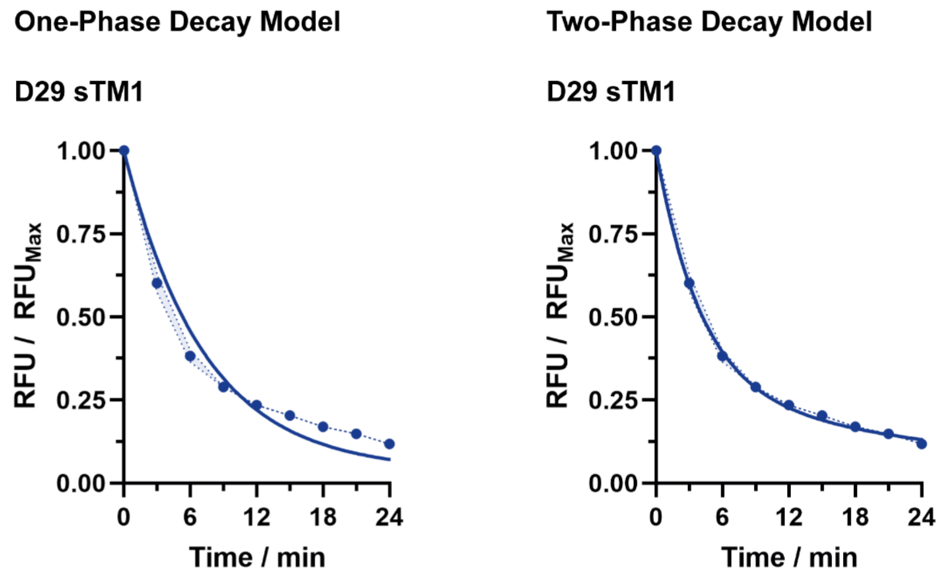

| | | |
|---|---|---|
| **R-Squared** | >0.966 | |
| **$T_{½}^{EDTA}$** | 5.0 ± 0.3 | 1.0 × |

| | | |
|---|---|---|
| **R-Squared** | >0.999 | |
| **$T_{½}^{EDTA}$ (Fast)** | 2.8 ± 0.1 | 0.72 × |
| **$T_{½}^{EDTA}$ (Slow)** | 20.6 ± 0.9 | 0.28 × |

**Figure S4:** Comparison of one-phase versus a two-phase exponential model used to fit the EDTA-dependent decay in the fluorescent signal of D29 sTM1. Notably, a two-phase model provides a better fit overall with the fast rate kinetic approximating the decay rates observed for the α-helically-locked D29 TM1 and D29 TM1$^{GP}$ variants.

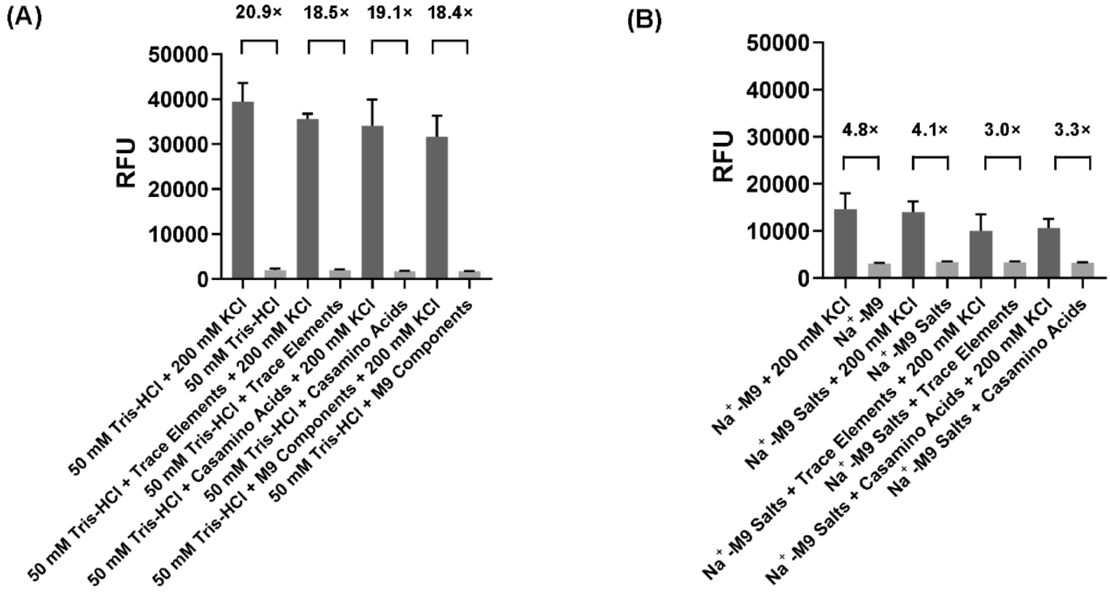

**Figure S5:** Analyzing the effect of medium and buffer composition on the response of GINKO2. The response of GINKO2 was measured with purified GINKO2 in the presence and absence of 200 mM KCl in **(A)** 50 mM Tris-HCl pH 8.0 and **(B)** $Na^+$-M9 medium supplemented with different additives as indicated (see **Tab. S2** for detailed composition): Trace Elements comprise 1x trace elements, $Na^+$-M9 comprises fully reconstituted $Na^+$-M9 medium for growing *E. coli*, $Na^+$-M9 Salts refers to the basic 1× $Na^+$-M9 stock solution, Casamino Acids are included at 0.1% (w/v).

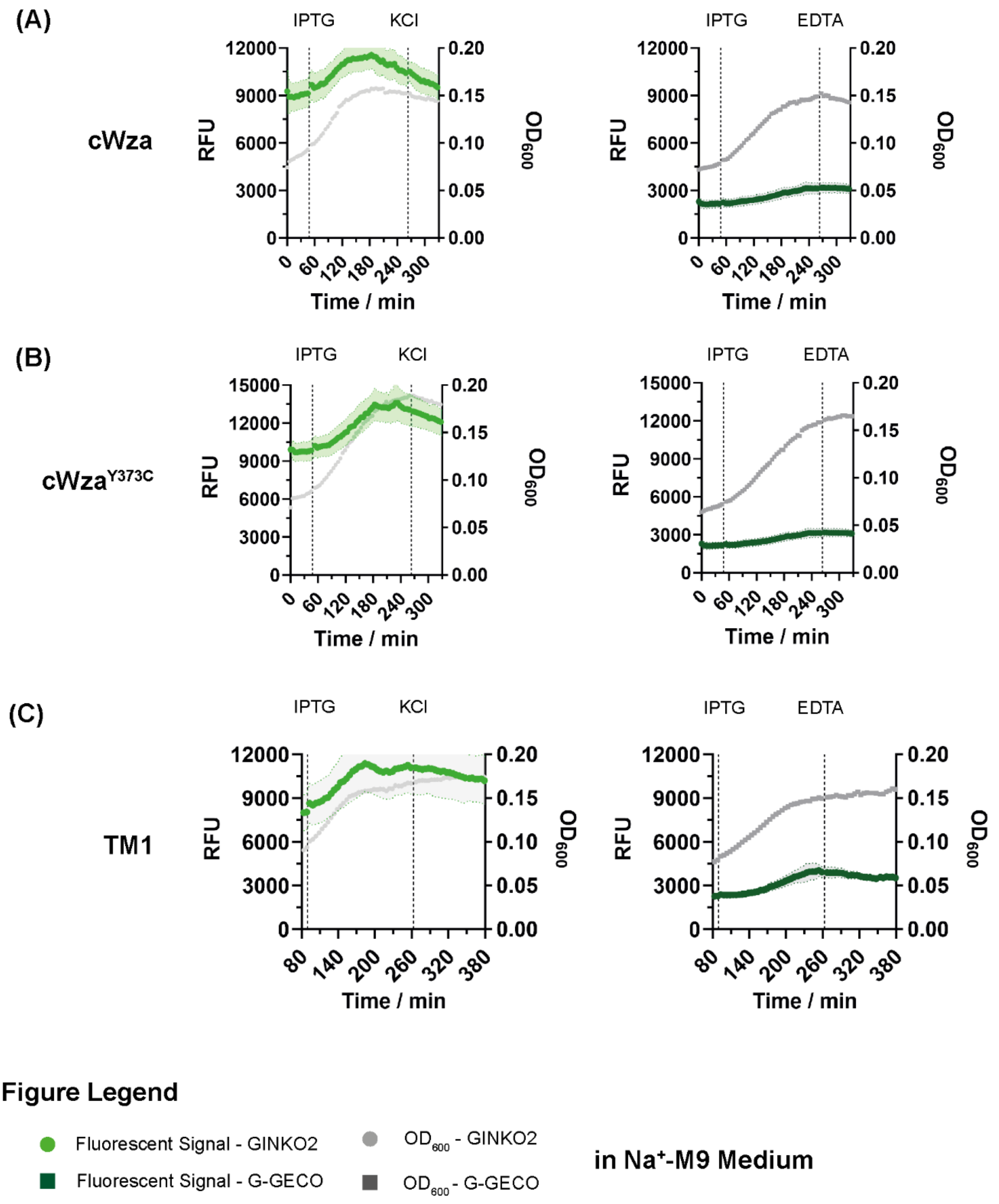

**Figure S6:** Summary of cWza, cWza$^{Y373C}$ and TM1 that did not reach an endpoint following growth of *E. coli* in $Na^+$-M9 medium. Nanopore assembly was monitored with $Ca^{2+}$-responsive G-GECO and the $K^+$-responsive GINKO2. The expression of individual nanopores was achieved under the control of the strong T7.100.WT.sRBS promoter. Nanopore expression was induced with 1 mM IPTG at the first time point indicated. Further, the $K^+$- and $Ca^{2+}$-dependent fluorescent signal was simultaneously quenched with 200 mM KCl and 5 mM EDTA at the second time point indicated.

**CCTM-LL-VI$_2$**

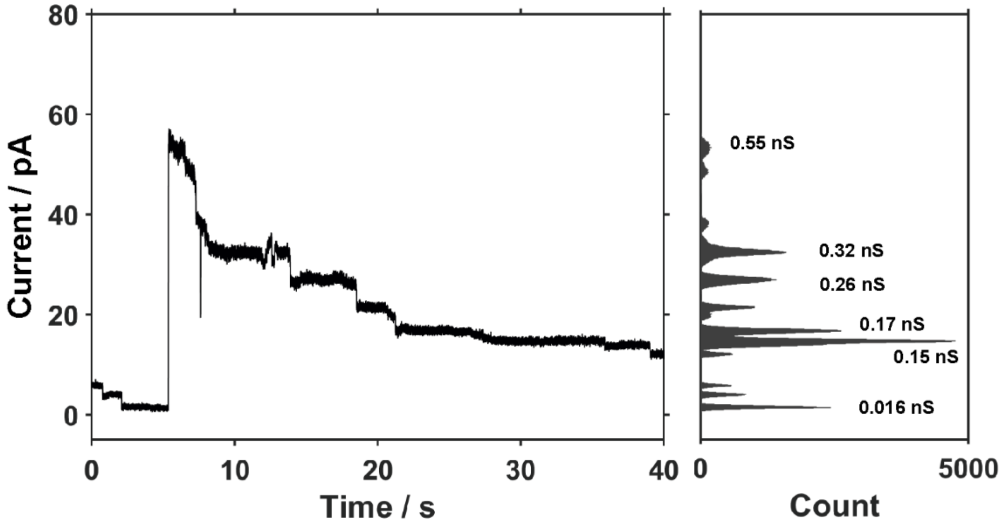

**Figure S7:** Additional electrophysiological current trace of the new recombinant CCTM-LL-VI$_2$ displays multiple heterogenous states with a steep initial increase which then levels between 0.26 – 0.32 nS before transitioning into a more uniform conductance of 0.15 – 0.17 nS. The analysis was conducted in 10 mM Tris-HCl, 500 mM $CaCl_2$ pH 8.0 at +100 mV.
